# Structure Bioinformatics of Eight Human ATP Synthase Fo Subunits and Their AlphaFold3-Predicted Water-Soluble QTY Analogs

**DOI:** 10.64898/2026.06.18.733091

**Authors:** Ziyu Wang, Edward Chen, Shuguang Zhang

## Abstract

Human mitochondrial ATP synthase is an essential rotary motor enzyme that produces most of the cellular ATP through oxidative phosphorylation. Its membrane-embedded Fo sector contains highly hydrophobic transmembrane subunits that are challenging to study in aqueous environments without detergents. This study explores whether applying the QTY code can reduce the hydrophobicity of selected ATP synthase Fo subunits while preserving their overall molecular structures. We applied the QTY code to eight human ATP synthase Fo subunits: ATP6, ATP8, ATPK, ATP68, ATPMK, AT5G1, AT5G2, and AT5G3. Hydrophobic amino acids leucine (L), isoleucine (I), valine (V), and phenylalanine (F) in transmembrane regions were systematically replaced with hydrophilic glutamine (Q), threonine (T), and tyrosine (Y). Four native subunits with available CryoEM structures from human ATP synthase (PDB: 8H9S) were superposed with their AlphaFold3-predicted QTY analogs. The native ATP synthase Fo subunits superposed well with their respective QTY analogs. For the CryoEM-native comparisons, RMSD values ranged from 0.565Å to 2.546Å. For the AlphaFold3-native comparisons of subunits without CryoEM structures, RMSD values ranged from 0.204Å to 0.297Å. Despite substantial QTY substitutions in the transmembrane regions, ranging from 38.89% to 50.79%, the QTY analogs retained similar overall folds, molecular weights, and isoelectric points. Hydrophobic surface analysis showed that the QTY analogs had reduced hydrophobic patches compared with their native counterparts, with average hydrophobicity decreasing from 0.2959 in native proteins to -1.1023 in QTY analogs. These structural bioinformatics studies suggest that the QTY code can be applied to ATP synthase Fo subunits to generate more hydrophilic, potentially water-soluble analogs while preserving overall structural similarity. These results extend the application of the QTY code to the membrane-embedded Fo sector of ATP synthase and provide a foundation for future experimental studies testing whether these QTY analogs can be expressed, purified, and evaluated for assembly or proton-transfer-related functions.

## Introduction

Mitochondrial ATP synthase is one of the most conserved and essential motor enzymes in the human body. It is the final step of oxidative phosphorylation in aerobic respiration, producing most of the adenosine triphosphate (ATP) that cells use as energy (1–4). ATP synthase plays key roles in both catalytic and transport mechanisms. Human ATP synthase is constructed with 29 subunits of 18 types, organized into membrane-intrinsic and membrane-extrinsic domains, linked by a central peripheral stalk. As such, the enzyme is a dual rotary motor built up of different subunits: the soluble catalytic F1 sector, where ATP is synthesized, and the membrane-embedded Fo sector, where proton translocation is converted into rotary motion.

The rotary mechanism comprises two elements: a rotor and a stator. The rotor includes the Fo c-ring and the central stalk, which rotates during proton translocation; the stator includes ATP6 and the peripheral stalk, which stabilize the catalytic F1 head against co-rotation (3,4).

The two sectors are coupled molecularly so that proton flow through the Fo sector drives conformational changes in F1 that synthesize ATP from adenosine diphosphate (ADP) and inorganic phosphate (1–5). Together, these sectors form the electrochemical proton gradient across the inner mitochondrial membrane and drive ATP production with great efficiency. Because ATP is a key step in cellular energy metabolism, even slight alterations to its structure could damage its assembly and coupling efficiency, causing severe physiological consequences and mitochondrial disease (1,3,6,7).

Within this complex, the Fo sector is significant for housing the proton-conducting motor and being the most hydrophobic part of the enzymes (2–5). In humans, the Fo sector is composed of the c-ring together with subunit ATP6, ATP8, and several associated membrane and peripheral stalk subunits to stabilize the rotor architecture (8–11). The Fo sector couples the movement of protons down their concentration gradient to the rotation of its c-ring, producing mechanical torque that is passed through the central stalk to the F1 sector. The Fo sector takes its name from its sensitivity to antibiotic oligomycin, which can bind to c-subunits to block proton flow, stopping rotation and ATP synthesis (9,12). The Fo sector is suitable for structural analysis, since it’s highly membrane-embedded, yet its functions rely on precise structural positioning, subunit interactions, and tight packing between hydrophobic helices (3,5,9,12).

Our study focuses on eight subunits, including: ATP6, ATP8, ATPK, ATP68, ATPMK, AT5G1, AT5G2, and AT5G3.

ATP6, encoded by mitochondrial DNA with gene MT-ATP6, is the central mechanical component among the Fo subunits in this study. It is highly hydrophobic, forming the core stator adjacent to the rotating c-ring and provides the offset proton half-channels that allow protons to enter and leave the membrane domain during catalysis (3–5,13). It helps generate the two offset proton half-channels required for proton entry, release, and generation of rotary movement (8,13). Many pathogenic analogs to MT-ATP6 can disrupt proton movement, a/c-ring coupling, or ATP synthesis rate (14–16).

ATP 8, encoded by mitochondrial DNA with gene MT-ATP8, is smaller and less involved in proton transfer, though studies have shown that it contributes to stabilization of ATP6 and the surrounding membrane domain during biogenesis of the complex. Interestingly, ATP6 and ATP8 are encoded in overlapping mitochondrial genes (9,11,17). Clinical studies have shown that MT-ATP8 analogs can contribute to mitochondrial diseases, though precise interpretation is often harder than MT-ATP6 because ATP8 is expressed at a lower protein level frequency, and because it appears to act more through structural support than direct catalysis (17).

ATP68, also known as subunit j or 6.8 kDa mitochondrial proteolipid protein (6.8PL), encoded by the gene ATP5MJ, is another Fo-associated subunit (9,10). Its function has been less defined historically, though recent studies have shown its use in locking ATP6 and ATP8 into their membrane assembly. It also helps dimerize the monomer complex via interactions between j subunits. Studies have shown that removal of ATP68 hindered cell growth and a partial loss of ATP6 and ATP8 (9,18).

ATPK, also known as subunit f, encoded by the gene ATP5MF, belongs to the supernumerary membrane region that helps maintain the structure of ATP synthase. Subunits like these occupy peripheral positions around the membrane motor, building up the membrane domain of the peripheral stalk. They contribute to local packing and maintain a wedge-shaped membrane, potentially helping with c-ring rotation (5,8–11). ATPK has been shown to assist in dimer stability, though its decrease has shown little effect on ATP synthetic capabilities or the amount of other subunits (19).

ATPMK, also known as subunit k, encoded by the gene ATP5MK, acts in a similar capacity as ATPK, appearing in the peripheral stalk of ATP synthase. It’s also been classified as a diabetes-associated protein in insulin-sensitive tissue (DAPIT) (5,8–11). Similar to ATPK, ATPMK promotes the structural stability of ATP synthase. Previous conclusions have classified ATPMK as a dispensable protein for the core functions of ATP synthase. However, recent studies have shown that suppression can cause a loss of ATP synthase population in the mitochondria and destabilization in the dimer structure. (20–22).

The three remaining subunits, AT5G1, AT5G2, and AT5G3, encoded by gene ATP5MC1, ATP5MC2, and ATP5MC3, respectively, are c-subunit isoforms that assemble into the c-ring in the Fo rotor. Subunit c is encoded by these three genes, though they only differ in their cleavable mitochondrial targeting peptides, resulting in generally identical mature proteins (13,23). These proteolipid subunits are the most hydrophobic among the Fo subunits in this study, together forming the rotating ring that converts proton movement into mechanical torque. The mechanism works as conserved acidic residues in the c-subunits bind and release protons during catalysis, resulting in a stepwise rotation relative to ATP6 that drives conformational changes through the central stalk into the F1 sector (4,8,13). Recent studies have connected mutations to multiple conditions, including cardiomyopathy and autosomal dominant spastic paraplegia and dystonia (24,25).

Taken together, these eight subunits span the major structural and functional classes within the Fo sector. ATP6 and AT5G c-subunits control proton translocation, ATP8 stabilizes the membrane domain during biogenesis, and ATP68, ATPK, and ATPMK represent smaller Fo subunits that contribute to assembly and maintaining structural integrity (5,8–11). Studying this set of subunits thus allows for evaluation of QTY analogs across mechanically and structurally important elements of the Fo motor.

ATP synthase Fo subunits are membrane-embedded and hydrophobic, making them challenging to study without the use of detergents. The QTY code solves this problem by converting hydrophobic transmembrane proteins into water-soluble analogs while seeking to preserve overall α-helix structure and function (26–28). In this approach, selected hydrophobic residues in transmembrane regions are substituted with structurally similar polar residues: leucine (L) is replaced by glutamine (Q), isoleucine (I) and valine (V) by threonine (T), and phenylalanine (F) by tyrosine (Y). These substitutions reduce the hydrophobic surface of membrane proteins, allowing for potential detergent-free studies while maintaining protein structure (26–29).

Recent advancements in cryogenic electron microscopy (CryoEM) have enabled high-resolution visualization of native membrane protein assemblies, such as ATP synthase, that have been previously difficult to stabilize (30–32). With these new developments, the molecular structure of human ATP synthase has been experimentally determined, split into three distinct states that represent different snapshots during its rotation, with roughly ∼120° between each state: State 1 (Protein Data Bank [PDB]: 8H9S) (33), state 2 (PDB: 8H9T) (34), and state 3, split into state 3a (PDB: 8H9U) (35) and state 3b (PDB: 8H9V) (36–38). For our study, we will be using state 1. However, these experimentally determined structures don’t contain all subunits, meaning we have to rely on computational tools like AlphaFold3 for full comparisons.

AlphaFold-based structure prediction has created opportunities to study membrane proteins in silico. After the release of AlphaFold2 in 2021, the QTY code has been applied to G protein-coupled receptors, glucose transporters, solute carrier transporters, ABC transporters, neurological transporters such as serotonin, dopamine, and norepinephrine transporters, and glutamate transporters (39–45). Reverse-QTY designs have also been used on human serum albumin for antitumor drug release and on the water-soluble CXCR4QTY receptor in a biomimetic sensing platform (46,47). AlphaFold3, released in 2024, extends structure predictions beyond single proteins and is now capable of predicting complex biomolecular interactions and behavior (48). Using AlphaFold3, the QTY code has also been applied to six human integral membrane enzymes and modeled ligand interactions (29).

Here, we present bioinformatic studies of four experimentally determined and four AlphaFold3-predicted human ATP synthase Fo subunits and their AlphaFold3-predicted water-soluble QTY analogs. We provide superpositions of the hydrophobic native subunits and their hydrophilic QTY analogs. We also provide the comparative hydrophobicity of molecular structures with their hydrophilic QTY analogs.

## Results and Discussion

### Rationale of the QTY Code on transmembrane helices

Integral transmembrane enzymes can be hard to study in aqueous solution because their transmembrane helices consist of hydrophobic residues essential for membrane localization. However, amino acids Q and L, T and V/I, and Y and F share similar electron density maps (26,27). The QTY code utilizes this similarly, and systematic replacements are made to reduce the hydrophobicity of integral transmembrane enzymes. The QTY code replaces hydrophobic amino acids with hydrophilic ones: leucine (L) with glutamine (Q), isoleucine (I) and valine (V) with threonine (T), and phenylalanine (F) with tyrosine (Y). Through these substitutions, the hydrophobic surface of the transmembrane regions is reduced without introducing major changes in residue size or helical packing tendency. Although sequence changes are introduced, QTY analogs remain similar to their native counterparts in isoelectric point (pI) and molecular weight (MW) (Figure 1).

**Figure 1.**
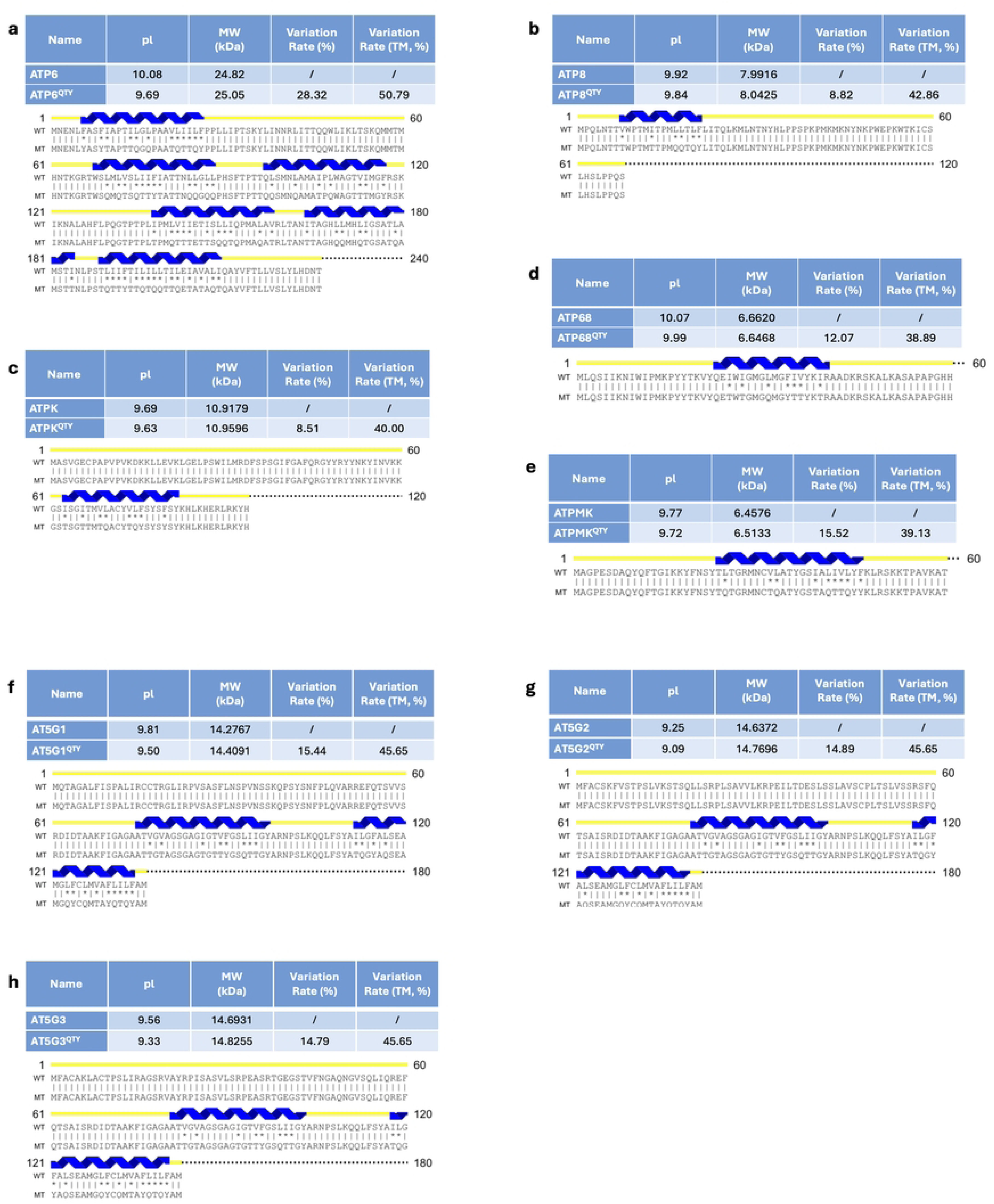
Protein sequence alignments of eight native ATP synthase Fo subunits with their water-soluble QTY analogs. The symbols | and * indicate whether amino acids are identical or different, respectively. Please note the Q, T, and Y amino acids replacing L, V and I, and F, respectively. The alpha helices (blue) are shown above the protein sequences. The characteristics of native and QTY analogs listed are isoelectric focusing point (pI), molecular weight (MW), total variation %, and transmembrane variation %. The alignments are: (a) ATP6 vs ATP6^QTY^, (b) ATP8 vs ATP8^QTY^, (c) ATPK vs ATPK^QTY^ (d) ATP68 vs ATP68^QTY^, (e) ATPMK vs ATPMK^QTY^, (f) AT5G1 vs AT5G1^QTY^, (g) AT5G2 vs AT5G2^QTY^, and (h) AT5G3 vs AT5G3^QTY^. Although there are substantial QTY changes in the transmembrane alpha helices (38.89–50.79%), the changes in MW and pI remain small.

### Protein sequence alignments and other characteristics

The amino acid sequences of ATP6, ATP8, ATPK, ATP68, ATPMK, AT5G1, AT5G2, and AT5G3 were aligned with their QTY analogs (Figure 1). Residues altered by the QTY code are marked by the “*” symbol. Overall, these substitutions led to amino acid changes ranging from 8.51% to 28.32% while the variation within the transmembrane domains ranged from 38.89% to 50.79% (Figure 1, Table 1). Nevertheless, the pI values remained largely unchanged, since Q, T, and Y are neutral amino acids that won’t introduce additional acidic or basic side chains. The MW values increased only slightly, which can be explained by the mass differences among the substituted residues: leucine (L, 131.17 Daltons) is lighter than glutamine (Q, 146.14 Daltons); isoleucine (I, 131.17 Daltons) is heavier whereas valine (V, 117.15 Daltons) is lighter than threonine (T, 119.12 Daltons); and phenylalanine (F, 165.19 Daltons) is lighter than tyrosine (Y, 181.19 Daltons). Additionally, the QTY substitutions replace carbon-rich hydrophobic side chains (Carbon, 12 Daltons) with oxygen and nitrogen bearing groups (-NH2, 14 Daltons and -OH, 16 Daltons), resulting in MW increases.

**Table 1.** Characteristics of eight human ATP synthase Fo subunits in this structural bioinformatic study. Native protein names, UniProt IDs, structural sources, tissue expression, and representative medical relevance are listed for ATP6, ATP8, ATPK, ATP68, ATPMK, AT5G1, AT5G2, and AT5G3. Tissue expression and medical relevance are representative and not exhaustive.

| Name (Uniprot ID) | Structure (Å, PDB ID) | Tissue expression | Medical relevance |
| --- | --- | --- | --- |
| ATP6 (P00846) | CryoEM (2.53 Å, 8H9S) | Ubiquitous | Mitochondrial disease |
| ATP8 (P03928) | CryoEM (2.53 Å, 8H9S) | Heart-enriched | Cardiomyopathy |
| ATPK (Q96IX5) | CryoEM (2.53 Å, 8H9S) | Ubiquitous | Complex V deficiency |
| ATP68 (P56378) | CryoEM (2.53 Å, 8H9S) | Skeletal muscle | ATP synthase assembly |
| AT5G1 (P05496) | AlphaFold3 predicted native structure | Ubiquitous | Oxidative phosphorylation |
| AT5G2 (Q06055) | AlphaFold3 predicted native structure | Ubiquitous | Oxidative phosphorylation |
| AT5G3 (P48201) | AlphaFold3 predicted native structure | Ubiquitous | Neuromuscular disease |
| ATPMK (Q96IX5) | AlphaFold3 predicted native structure | Ubiquitous | Complex V deficiency |

### AlphaFold3 Predictions and Model Confidence

Structures of the QTY analogs and subunits without CryoEM structures were predicted using AlphaFold3 through the online AlphaFold server (48,49). Model quality was assessed using the confidence metrics, including Local Distance Difference Test (pLDDT), predicted Template Modeling score (pTM), and Predicted Aligned Error (PAE)

The pLDDT measures the model’s confidence in each residue (48,50). In AlphaFold3 predictions, dark blue indicates high confidence (pLDDT > 90), light blue indicates good confidence (pLDDT between 70 and 90), yellow indicates moderate confidence (pLDDT between 50 and 70), and orange indicates low confidence (pLDDT < 50) (48). In our predictions, most transmembrane regions were dark or light blue, indicating well-ordered helical structures, whereas terminal regions more often appeared yellow or orange, indicating greater flexibility or disorder (Figure 2) (51). Whenever AlphaFold3 was used in a structural comparison, we excluded low-confidence and unstructured regions from both the predicted structure and the corresponding structure before analysis, so that the comparison focused on well-resolved and meaningful differences. The pTM measures overall model confidence by comparing protein structures with known templates (51). In our study, the predicted QTY analogs achieved an average pTM score of 0.732, indicating good overall structural reliability (Figure 2). PAE evaluates confidence in the relative positioning of residue pairs within the structure model (52). In our predictions, the PAE plots were largely dominated by low-error regions, indicating high confidence in the relative spatial arrangement of most residues (Figure 2). Together, the pLDDT, pTM, and PAE results support the overall reliability of AlphaFold3 models for structural analysis of the QTY analogs and non-CryoEM structures. Nevertheless, these models remain computational predictions and will require experimental validation to confirm their structures.

**Figure 2.**
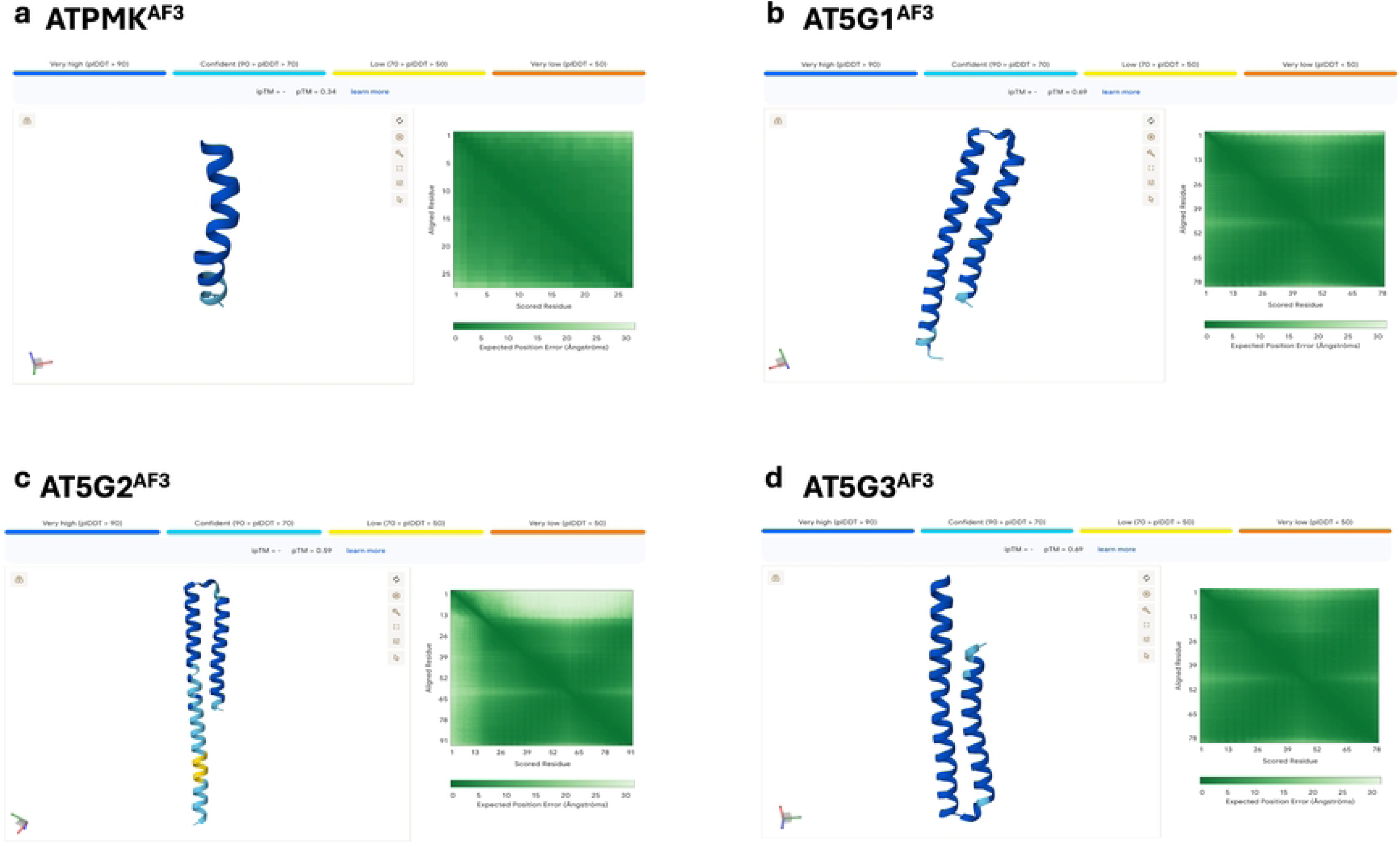
AlphaFold3 confidence summaries for ATP synthase Fo subunits without CryoEM structures. AlphaFold3-predicted native structures for **a)** ATPMK^AF3^, **b)** AT5G1^AF3^, **c)** AT5G2^AF3^, and **d)** AT5G3^AF3^ are shown with per-residue confidence (pLDDT; blue = high confidence, yellow/orange/red = lower confidence), predicted aligned error (PAE) heatmaps, and predicted template modeling (pTM) scores. These AlphaFold3-predicted native structures were used as reference native models for native vs QTY structural superpositions and hydrophobicity surface comparisons in Figures x—y. Protein sequences of unstructured loops were removed for clarity.

### Reference Native Structure Selection

Not all subunits had complete experimental native CryoEM structures. Thus, ATP6, ATP8, ATPK, and ATP68 were compared using native CryoEM structures, while AT5G1, AT5G2, AT5G3, and ATPMK were compared using AlphaFold3-predicted native models. This ensured more consistent full-length protein comparisons between native and QTY structures. Further explanations in the Methods section.

### Superpositions of select CryoEM-determined native ATP synthase Fo subunits and their water-soluble QTY analogs

We ask whether the molecular structure of four ATP synthase Fo transmembrane subunits can be superposed with their respective QTY analogs (Figure 3). The native structures of ATP6, ATP8, ATPK, and ATP68 (PDB: 8H9S) have been experimentally determined via CryoEM, whereas the QTY analogs were predicted via AlphaFold3. The superpositions are: ATP6^CryoEM^ *vs* ATP6^QTY_AF3^, ATP8^CryoEM^ *vs* ATP8^QTY_AF3^, ATPK^CryoEM^ *vs* ATPK^QTY_AF3^, and ATP68^CryoEM^ *vs* ATP68^QTY_AF3^.

**Figure 3.**
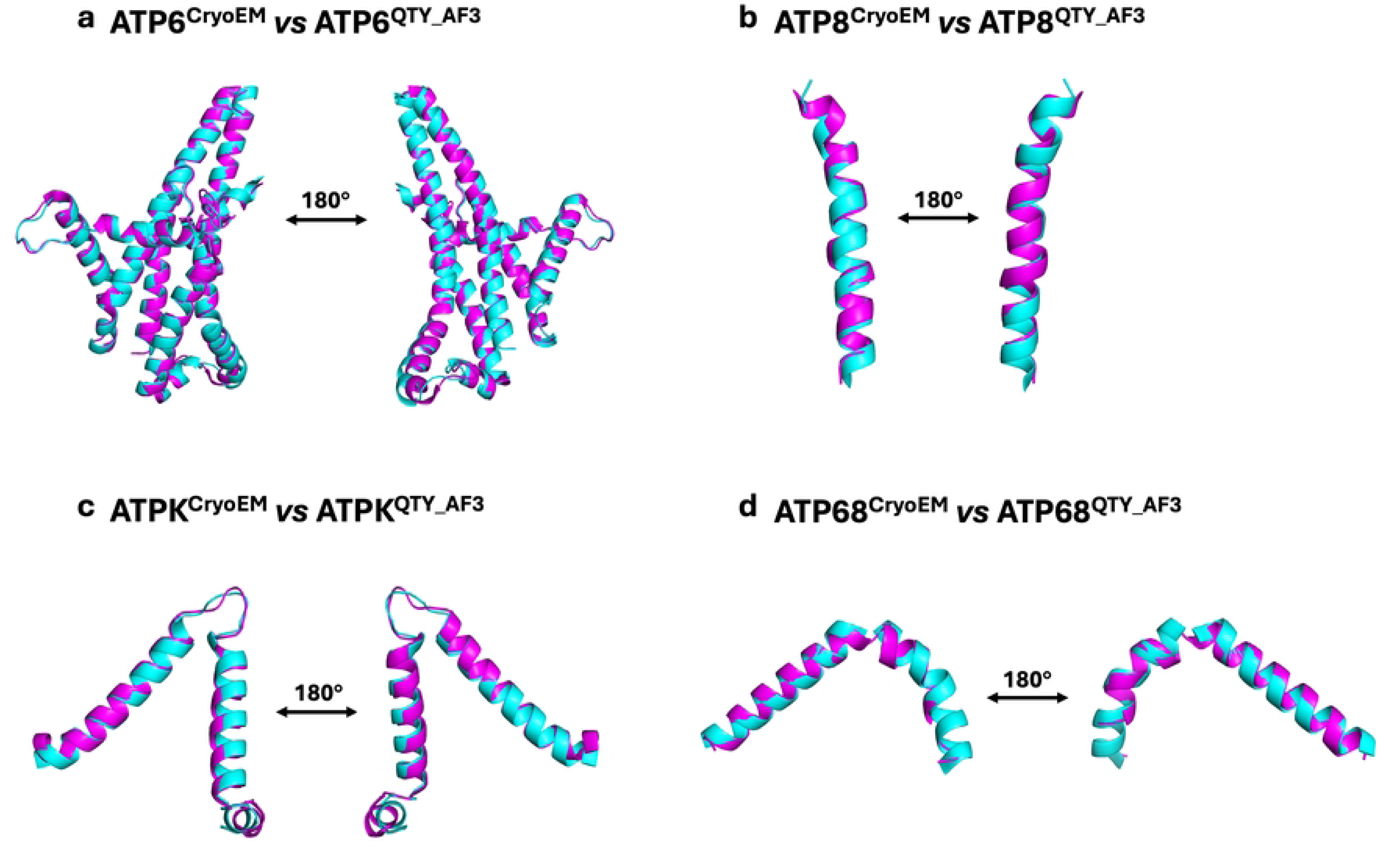
Superpositions of select CryoEM-determined native ATP synthase Fo subunits and their water-soluble QTY analogs. The CryoEM-determined structures of the native subunits are obtained from the Protein Data Bank (PDB). CryoEM structures (magenta) are superposed with their QTY analogs (cyan) predicted by AlphaFold3. These superpositions show that the native subunits share similar structure to their QTY analogs. Unstructured loops with low confidence were removed. Additionally, because the native ATP68 CryoEM structure is bent, ATP68 was analyzed using segment-based superposition from residues 2-20 and 21-42. Both RMSDs are reported in their respective order. Two views are shown for each protein (180°). Panels show: **a)** ATP6^CryoEM^ *vs* ATP6^QTY_AF3^ (RMSD = 0.565 Å), **b)** ATP8^CryoEM^ *vs* ATP8^QTY_AF3^ (RMSD = 2.546 Å), **c)** ATPK^CryoEM^ *vs* ATPK^QTY_AF3^ (RMSD = 1.180 Å), and **d)** ATP68^CryoEM^ *vs* ATP68^QTY_AF3^ (RMSD = 1.608 Å, 0.790 Å).

The native CryoEM structures and their AlphaFold3-predicted QTY analogs superposed well. The root-mean-square deviation (RMSD) values ranged from 0.565 Å to 2.546 Å (Figure 3, Table 2), specifically: a) ATP6^CryoEM^ *vs* ATP6^QTY_AF3^ (RMSD = 0.565 Å), b) ATP8^CryoEM^ *vs* ATP8^QTY_AF3^ (RMSD = 2.546 Å), c) ATPK^CryoEM^ *vs* ATPK^QTY_AF3^ (RMSD = 1.180 Å), and d) ATP68^CryoEM^ *vs* ATP68^QTY_AF3^ (RMSD = 1.608 Å, 0.790 Å). As seen in Figure 3, these structures share similar overall folds despite a 38.89–50.79% of replacements in transmembrane region amino acids using the QTY code.

**Table 2.** Protein characteristics of eight human ATP synthase Fo subunits and their QTY analogs. Abbreviations: pI, isoelectric point; MW, molecular weight; TM, transmembrane; AF3, AlphaFold3; CryoEM, cryo-electron microscopy; –, not applicable; RMSD, root-mean-square deviation. The eight ATP synthase Fo subunits are listed in the same order as Figures 1 and 2. Native structures for ATP6, ATP8, ATPK, and ATP68 were derived from the CryoEM human ATP synthase structure (PDB: 8H9S), whereas native structures for ATPMK, AT5G1, AT5G2, and AT5G3 were predicted by AlphaFold3. RMSDs were calculated after truncating unresolved and low-confidence regions in native proteins and corresponding residues in QTY analogs. ATP68 has two RMSD values reported because the protein was superposed in two segments to account for the bend observed in the CryoEM structure. QTY substitutions within TM regions are substantial, ranging from 38.89% to 50.79%, while overall sequence changes range from 8.51% to 28.32%.

| Name | RMSD (Å) | pI | Mw (kDa) | TM Variation (%) | Overall Variation (%) |
| --- | --- | --- | --- | --- | --- |
| ATP6<sup>CryoEM</sup> | - | 10.08 | 24.82 | - | - |
| ATP6<sup>QTY_AF3</sup> | 0.565 | 9.69 | 25.05 | 50.79 | 28.32 |
| ATP8<sup>CryoEM</sup> | - | 9.92 | 7.99 | - | - |
| ATP8<sup>QTY_AF3</sup> | 2.546 | 9.84 | 8.04 | 42.86 | 8.82 |
| ATPK<sup>CryoEM</sup> | - | 9.69 | 10.92 | - | - |
| ATPK<sup>QTY_AF3</sup> | 1.180 | 9.63 | 10.96 | 40.00 | 8.51 |
| ATP68<sup>CryoEM</sup> | - | 10.07 | 6.66 | - | - |
| ATP68<sup>QTY_AF3</sup> | 1.608, 0.790 | 9.99 | 6.65 | 38.89 | 12.07 |
| AT5G1<sup>AF3</sup> | - | 9.81 | 14.28 | - | - |
| AT5G1<sup>QTY_AF3</sup> | 0.252 | 9.50 | 14.41 | 45.65 | 15.44 |
| AT5G2<sup>AF3</sup> | - | 9.25 | 14.64 | - | - |
| AT5G2<sup>QTY_AF3</sup> | 0.271 | 9.09 | 14.77 | 45.65 | 14.89 |
| AT5G3<sup>AF3</sup> | - | 9.56 | 14.69 | - | - |
| AT5G3<sup>QTY_AF3</sup> | 0.297 | 9.33 | 14.82 | 45.65 | 14.79 |
| ATPMK<sup>AF3</sup> | - | 9.77 | 6.46 | - | - |
| ATPMK<sup>QTY_AF3</sup> | 0.204 | 9.72 | 6.51 | 39.13 | 15.52 |

### Superpositions of select AlphaFold3-predicted ATP synthase Fo subunits with experimental data and their water-soluble QTY analogs

Although these four ATP synthase Fo subunits have experimentally determined CryoEM native structures available, we performed an additional test by superposing their AlphaFold3-predicted native structures with their AlphaFold3-predicted water-soluble QTY analogs (Figure 4). Both the native structure and QTY analogs were predicted via AlphaFold3. The structures superposed well with low RMSD, specifically: a) ATP6^AF3^ *vs* ATP6^QTY_AF3^ (RMSD = 0.341 Å), b) ATP8^AF3^ *vs* ATP8^QTY_AF3^ (RMSD = 0.238 Å), c) ATPK^AF3^ *vs* ATPK^QTY_AF3^ (RMSD = 2.583 Å), and d) ATP68^AF3^ *vs* ATP68^QTY_AF3^ (RMSD = 0.955 Å, 0.384 Å). These superpositions further support that water-soluble QTY analogs share very similar structures with the native ATP synthase Fo subunits, and it also validates AlphaFold3’s capability and accuracy.

**Figure 4.**
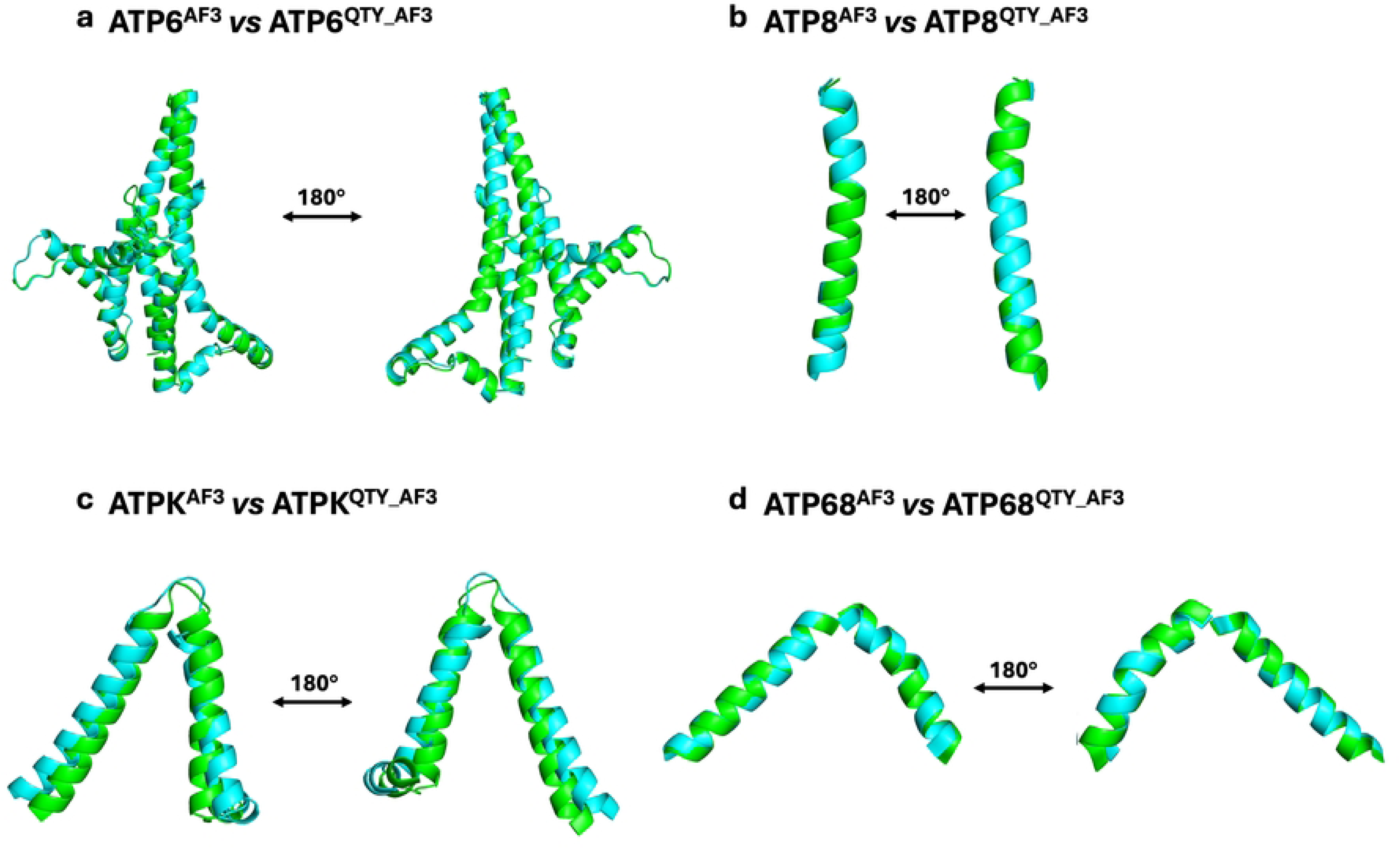
Superpositions of select AlphaFold3-Predicted native ATP synthase Fo subunits with experimental data and their water-soluble QTY analogs. AlphaFold3-predicted native structures (green) are superposed with their corresponding QTY analogs (cyan). Unstructured loops with low confidence were removed. Two views are shown for each protein (180°). RMSDs for both halves of ATP68 are reported in their respective order. Two views are shown for each protein (180°). Panels show: **a)** ATP6^AF3^ *vs* ATP6^QTY_AF3^ (RMSD = 0.341 Å), **b)** ATP8^AF3^ *vs* ATP8^QTY_AF3^ (RMSD = 0.238 Å), **c)** ATPK^AF3^ *vs* ATPK^QTY_AF3^ (RMSD = 2.583 Å), and **d)** ATP68^AF3^ *vs* ATP68^QTY_AF3^ (RMSD = 0.955 Å, 0.384 Å).

### Superpositions of select CryoEM-determined native ATP synthase Fo subunits with their AlphaFold3-predicted native structures and water-soluble QTY analogs

For ATP synthase Fo subunits with available CryoEM structures, we superposed i) all available CryoEM-determined native ATP synthase Fo subunits ii) AlphaFold3-predicted native ATP synthase Fo subunits and iii) AlphaFold3-predicted QTY analogs. The three different structures superposed well (Figure 5). This validates that there is a high similarity between experimentally determined structures and their QTY analogs, and between AlphaFold3-predicted native structures and QTY analogs (Figures 1 and 3). We also saw the high similarity between CryoEM structures and AlphaFold3-predicted native structures, providing a rationale for our subsequent comparisons using AlphaFold3-predicted native structures as substitutions of CryoEM structures when experimental data weren’t available. This suggests that the water-soluble QTY analogs could be expressed, purified, and used as soluble antigens to generate therapeutic monoclonal antibodies.

**Figure 5.**
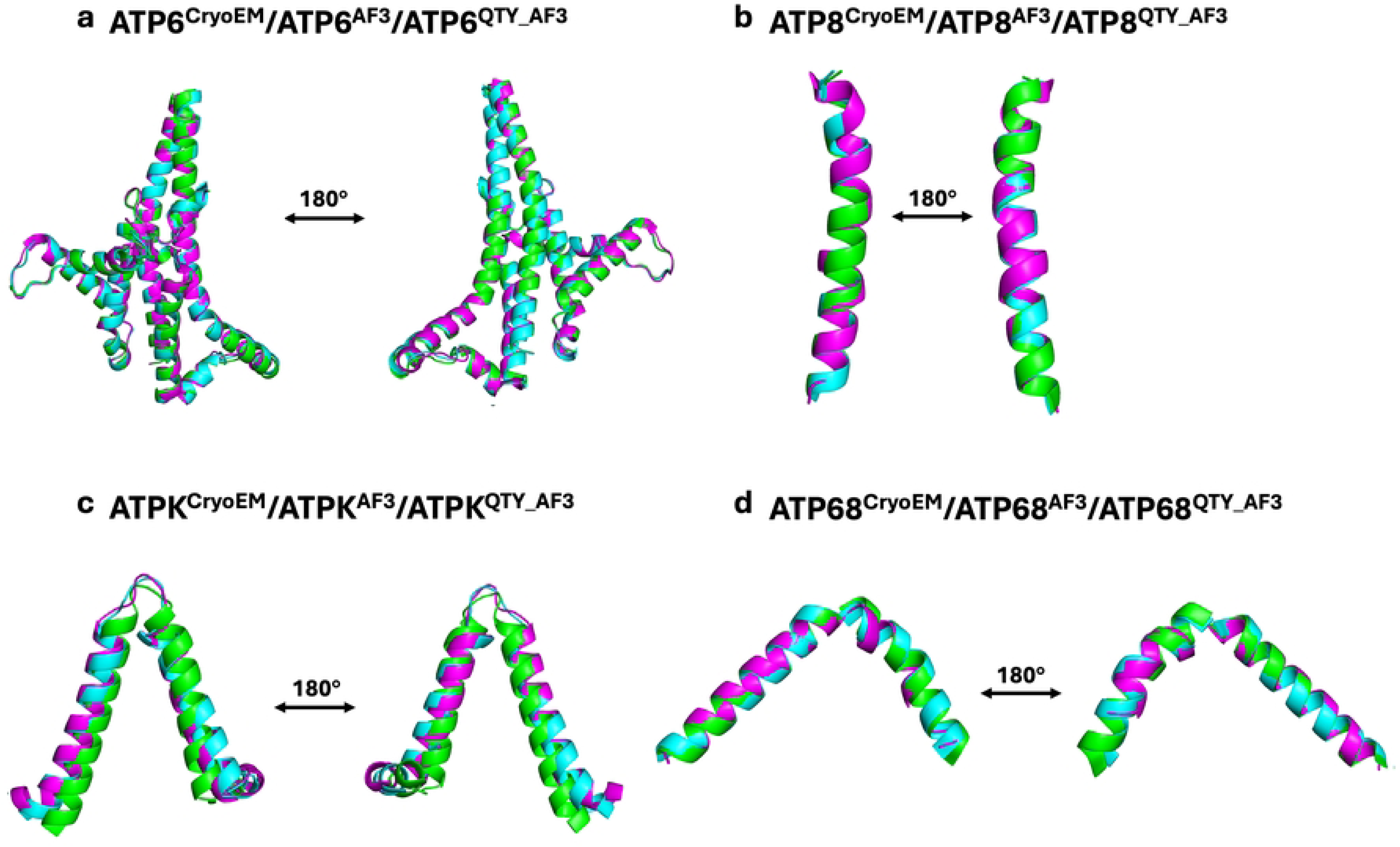
Superpositions of select CryoEM-determined native ATP synthase Fo subunits with their AlphaFold3-predicted native structures and water-soluble QTY analogs. Superposition of i) all available CryoEM-determined native ATP synthase Fo subunits (magenta) with ii) select AlphaFold3-predicted native ATP synthase Fo subunits (green) and iii) select AlphaFold3-predicted water-soluble QTY analogs (cyan). Individual superpositions are shown in Figures 4 and 3. All three structures superposed well visually, with insignificant differences shown. Two views are shown for each protein (180°). Panels show: **a)** ATP6^CryoEM^/ATP6^AF3^/ATP6^QTY_AF3^, **b)** ATP8^CryoEM^/ATP8^AF3^/ATP8^QTY_AF3^, **c)** ATPK^CryoEM^/ATPK^AF3^/ATPK^QTY_AF3^, and **d)** ATP68^CryoEM^/ATP68^AF3^/ATP68^QTY_AF3^.

### Superpositions of select AlphaFold3-predicted native ATP synthase Fo subunits without experimental data and their water-soluble QTY analogs

Since some subunits lacked CryoEM structures, and we’ve already shown the validity of AlphaFold3-predicted native structures in substitution for experimentally determined structures, we ask whether the AlphaFold3-predicted molecular structure of four ATP synthase Fo transmembrane subunits can be superposed with their respective QTY analogs (Figure 6). The native structures that lacked experimental data of AT5G1, AT5G2, AT5G3, and ATPMK were predicted via AlphaFold3, and their QTY analogs were also predicted via AlphaFold3. The superpositions are: AT5G1^AF3^ *vs* AT5G1^QTY_AF3^, AT5G2^AF3^ *vs* AT5G2^QTY_AF3^, AT5G3^AF3^ *vs* AT5G3^QTY_AF3^, and ATPMK^AF3^ *vs* ATPMK^QTY_AF3^.

**Figure 6.**
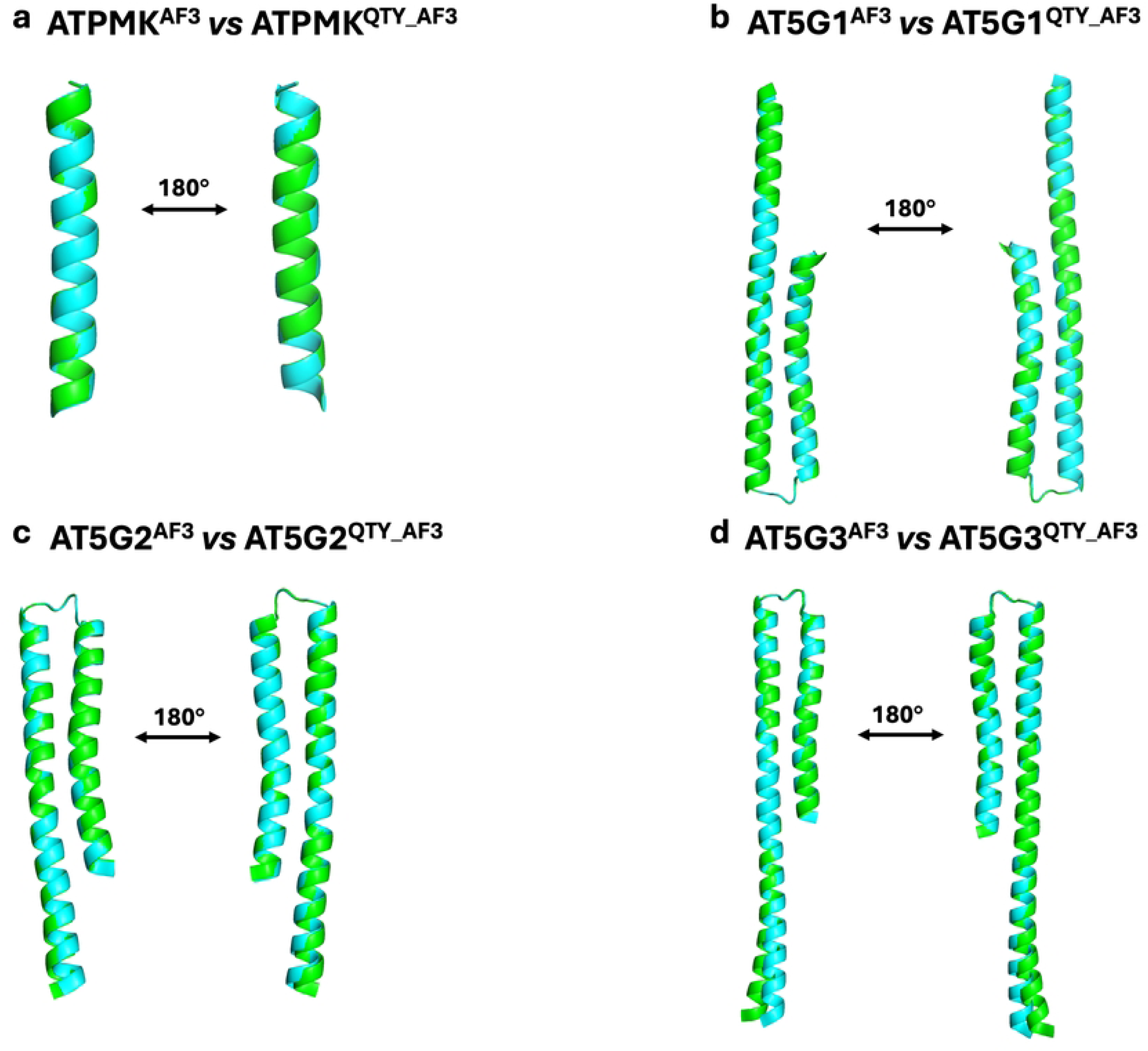
Superpositions of select AlphaFold3-predicted native ATP synthase Fo subunits without experimental data and their water-soluble QTY analogs. AlphaFold3-predicted native structures (green) are superposed with their corresponding QTY analogs (cyan). Unstructured loops with low confidence were removed. Two views are shown for each protein (180°). Panels show: **a)** ATPMK^AF3^ *vs* ATPMK^QTY_AF3^ (RMSD = 0.204 Å), **b)** AT5G1^AF3^ *vs* AT5G1^QTY_AF3^ (RMSD = 0.252 Å), **c)** AT5G2^AF3^ *vs* AT5G2^QTY_AF3^ (RMSD = 0.271 Å), and **d)** AT5G3^AF3^ *vs* AT5G3^QTY_AF3^ (RMSD = 0.297 Å).

The AlphaFold3-predicted native structures and their AlphaFold3-predicted QTY analogs superposed well. The RMSD values ranged from 0.204 Å to 0.297 Å (Figure 6, Table 2), specifically: a) AT5G1^AF3^ *vs* AT5G1^QTY_AF3^ (RMSD = 0.252 Å), b) AT5G2^AF3^ *vs* AT5G2^QTY_AF3^ (RMSD = 0.271 Å), c) AT5G3^AF3^ *vs* AT5G3^QTY_AF3^ (RMSD = 0.297 Å), and d) ATPMK^AF3^ *vs* ATPMK^QTY_AF3^ (RMSD = 0.204 Å). As seen in Figure 6, these structures share similar overall folds despite a 39.13–45.65% of replacements in transmembrane region amino acids using the QTY code.

### Analysis of the hydrophobic surface of CryoEM-determined and AlphaFold3-predicted native ATP synthase Fo subunits and their water-soluble QTY analogs

The eight ATP synthase Fo transmembrane subunits selected are inherently hydrophobic and water-insoluble, especially in the transmembrane alpha helical domains. The eight proteins have an average hydrophobicity of 0.2959 in their transmembrane regions. Typically, for protein studies, detergents are used to solubilize the native proteins after removing them from the lipid bilayer. Without proper tools, these proteins can aggregate, precipitate, or denature in aqueous solutions.

Hydrophobic surface areas are highlighted in yellow (Figures 7, 8), embedded in the hydrophobic lipid bilayer. Here, Leucine (L), Isoleucine (I), Valine (V), and Phenylalanine (F) interact with lipid molecules to exclude water, leading to a hydrophobic and nonpolar patch.

**Figure 7.**
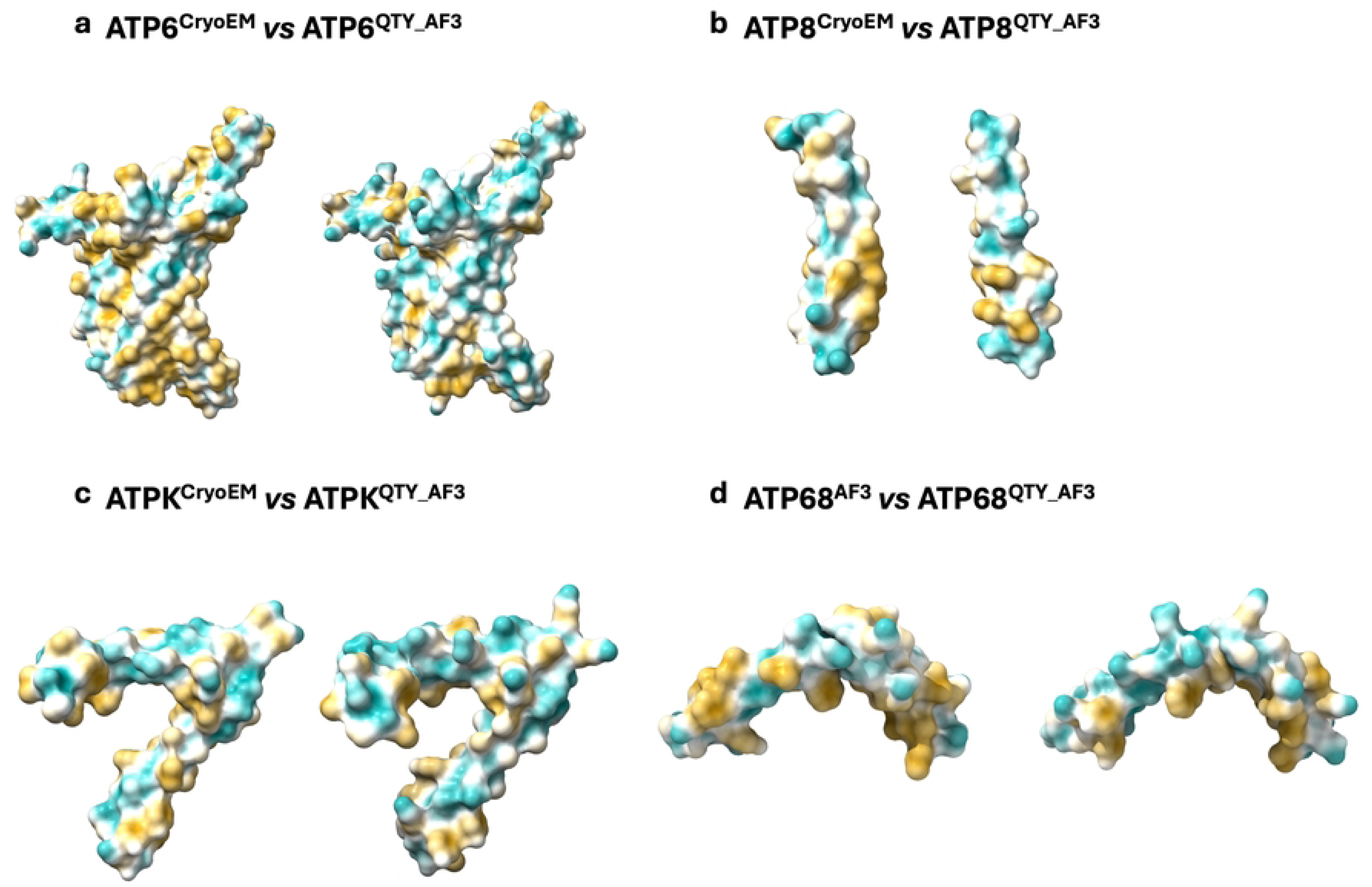
Hydrophobic surface comparison of CryoEM-determined ATP synthase Fo subunits and AlphaFold3-predicted QTY analogs. Molecular surface hydrophobicity is mapped as hydrophobic (yellow) and hydrophilic (cyan). The native subunits have many hydrophobic residues L, I, V, and F in their transmembrane helices. QTY analogs were generated by substituting L → Q, I/V → T, and F → Y to reduce transmembrane hydrophobicity. Unstructured loops with low confidence were removed. Two views are shown for each protein (180°). Panels show: **a)** ATP6^CryoEM^ *vs* ATP6^QTY_AF3^, **b)** ATP8^CryoEM^ *vs* ATP8^QTY_AF3^, **c)** ATPK^CryoEM^ *vs* ATPK^QTY_AF3^, and **d)** ATP68^CryoEM^ *vs* ATP68^QTY_AF3^.

**Figure 8.**
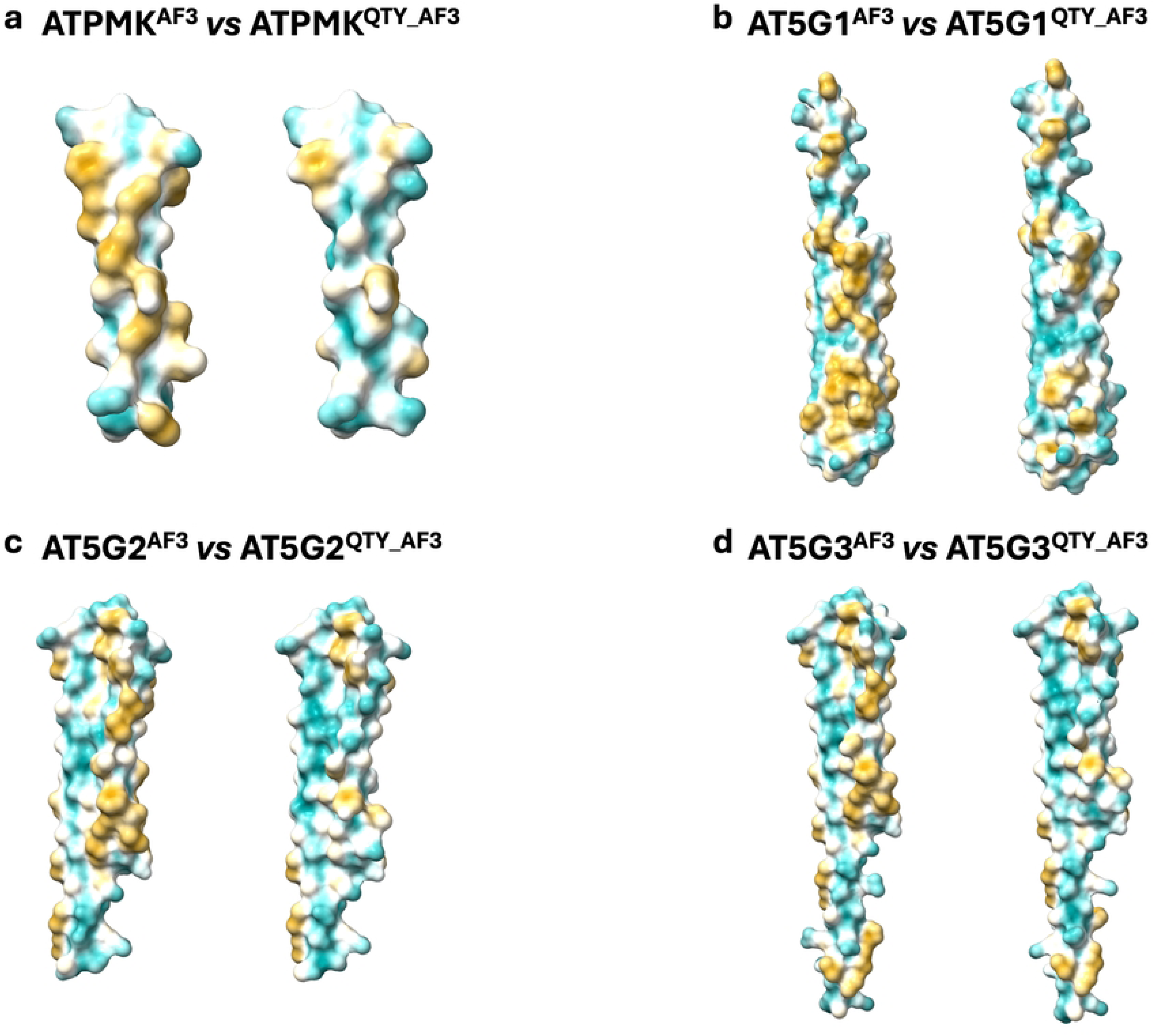
Hydrophobic surface comparison of AlphaFold3-predicted native ATP synthase Fo subunits and their QTY analogs. Molecular surface hydrophobicity is mapped as hydrophobic (yellow) and hydrophilic (cyan). The native subunits have many hydrophobic residues L, I, V, and F in their transmembrane helices. QTY analogs were generated by substituting L → Q, I/V → T, and F → Y to reduce transmembrane hydrophobicity. Unstructured loops with low confidence were removed. Two views are shown for each protein (180°). Panels show: **a)** ATPMK^AF3^ *vs* ATPMK^QTY_AF3^, **b)** AT5G1^AF3^ *vs* AT5G1^QTY_AF3^, **c)** AT5G2^AF3^ *vs* AT5G2^QTY_AF3^, and **d)** AT5G3^AF3^ *vs* AT5G3^QTY_AF3^.

By applying the QTY code, we systematically replaced hydrophobic amino acids L, I/V, and F, with hydrophilic amino acids Glutamine (Q), Threonine (T), and Tyrosine (Y), reducing hydrophobic surfaces. The average hydrophobicity of the eight proteins decreased from 0.2959 in their native analogs to -1.1023 in the QTY analogs. The decrease in hydrophobicity was significant, and the QTY code maintained overall alpha helical structure consistently. This is consistent with previous experiments, where QTY analogs of chemokine and cytokine receptors were shown to retain structural integrity, stability, ligand-binding ability, and become water-soluble.

### Rationale for Selecting ATP Synthase Fo Subunits for Our Study

The QTY code can be applied to a wide range of membrane enzymes, such as initial studies focusing on a total of seven chemokine receptors and four cytokine receptors. After carrying out a wide range of biophysical, biochemical, and molecular experiments (27,28,45). This study expands on the demonstrated success of the QTY code, selecting eight subunits from the ATP Synthase Fo: ATP6, ATP8, ATPK, ATP68, ATPMK, AT5G1, AT5G2, and AT5G3. These proteins were chosen because they were integral membrane components of the mitochondrial ATP complex, an extremely conserved, crucial, and complex molecular machine responsible for proton translocation across the membrane and ATP production, producing most of the energy for a cell. As membrane-associated Fo subunits, these proteins are valuable in providing a test case for whether the QTY code can expand beyond previously-studied receptors and transporters to cases involved in bioenergetic membrane machinery.

These proteins also hold significant biological and medical relevance. ATP synthase is a key component for oxidative phosphorylation, part of cellular respiration, and defects in its membrane sector have been associated with impaired mitochondrial energy metabolism, severe neuromuscular diseases like Parkinson’s, Alzheimer’s, and Amyotrophic Lateral Sclerosis, and other diseases like fragile X syndrome, the fatal neurodegenerative Batten disease, and cardiovascular complications (53–55). Additionally, the strongly hydrophobic nature of transmembrane alpha helices in Fo subunits makes them hard to study experimentally in aqueous solutions without detergents. This combination of functional importance and structural architecture makes the ATP synthase Fo sector a suitable system for testing whether QTY substitutions can still reduce hydrophobicity while preserving overall fold and enabling informative structural comparisons across a new type of protein system.

### Summary

In our current study, we applied the QTY code to eight ATP Synthase Fo subunits: ATP6, ATP8, ATPK, ATP68, ATPMK, AT5G1, AT5G2, and AT5G3. We generated corresponding water-soluble QTY analogs, comparing them with CryoEM or AlphaFold3-generated native structures for bioinformatics analysis. We employed numerous in silico computational tools to analyze the sequences and structural characteristics. Although QTY substitutions introduced changes in amino acid sequence and composition, the native and QTY proteins remained similar in key properties, including isoelectric point and molecular weight. Overall fold patterns and molecular integrity were preserved, suggesting that QTY analogs retain their structural functionality. QTY analogs also showed significantly reduced hydrophobic surfaces, further suggesting that the QTY code is a good approach to model water-soluble analogs of ATP Synthase Fo subunits. These results are consistent with previous studies of the QTY code, which have focused on chemokine receptors, cytokine receptors, ABC transporters, rhomboid proteases, and other integral membrane proteins (27,28,39–45).

The ATP synthase Fo sector is an essential and meaningful system in testing broader applications of the QTY code. These subunits form a membrane-embedded proton-translocating system for ATP synthase, a highly conserved and essential component of mitochondrial ATP production as part of oxidative phosphorylation during cellular respiration. Defects have been associated with a wide spectrum of human disease, including but not limited to neurodegenerative, neuromuscular, metabolic, and cardiovascular disorders (53–55). For this reason, the ability to develop more hydrophobic yet structurally similar Fo subunit analogs becomes a crucial step for future biochemical characterization and potential development of soluble antigens or disease-relevant membrane components (27,28,39,45).

Future studies could expand by testing whether ATP synthase Fo subunits’ QTY analogs preserve the capability to interact with other subunits and conduct proton-transfer related features required for assembly and function. ATP synthase comprises the membrane-embedded Fo rotor and F1 catalyst in the matrix, both requiring intimate and proper protein binding to function. Thus, ATP synthase function, specifically the Fo rotor, depends on interactions between multiple subunits. For example, ATP8, included in this study, interacts with subunit b, which was not included. In the native enzyme, ATP8 stabilizes the association of ATP6 with the ATP synthase peripheral stalk through binding with subunit b. Meanwhile, proton translocation occurs at the ATP6–c-ring interface. ATP6 and the c-subunit homologs AT5G1, AT5G2, and AT5G3 were included in this study, but further studies should evaluate these subunits as assembled proton-conducting interfaces (9–11,13). In AlphaFold3, this could be approached by modeling multimeric Fo sector complexes to test whether native-like interactions remain after QTY substitution (48).

Because the study is purely computational, the current conclusions should be interpreted within the scope of in silico structural bioinformatic experiments. AlphaFold3 confidence metrics support its reliability, but experimental validation ought to be performed to determine whether ATP Synthase Fo subunits’ QTY analogs can be expressed as functioning water-soluble proteins (48). Nevertheless, current results indicate that the QTY code can be extended beyond previously studied membrane enzymes to the ATP Synthase Fo sector, an important group of bioenergetic membrane proteins with great biological and medical importance.

## Conclusion

Nature has produced three chemically distinct yet structurally similar alpha-helices: 1) Type I hydrophilic alpha-helices, occurring in water-soluble proteins like hemoglobin, lysosome, growth factors, cytokines, and antibodies; 2) Type II hydrophobic alpha-helices, occurring in integral membrane proteins like G-protein coupled receptors, transporters, ion channels, and photosynthetic electron transport complexes; and 3) Type III amphiphilic alpha-helices, containing both hydrophobic and hydrophilic sides. These three Types of helices remain structurally highly similar in core geometry, despite substantial protein sequence differences in amino acid makeup and physicochemical properties. This principle is the molecular basis for the QTY code (27). The QTY code is a very simple protein editing tool that can directly edit Type II hydrophobic helices of membrane protein to their hydrophilic Type I water-soluble helices. Thus, the water-soluble QTY analogs can be used for a variety studies beyond the native membrane proteins.

## Methods

### Protein sequence alignments and other characteristics

We obtained the native sequence for eight human proteins using UniProt (https://www.uniprot.org/): ATP6, ATP8, ATPK, ATP68, ATPMK, AT5G1, AT5G2, and AT5G3. UniProt transmembrane-region annotations were used for the transmembrane regions and visualized as topology diagrams with Protter (https://wlab.ethz.ch/protter/). Water-soluble QTY analogs were generated by applying the QTY code to native FASTA sequences using the PSS QTY website (https://pss.sjtu.edu.cn/) (56). The server also produced several other protein characteristics (e.g. MWs, pI values, hydrophobicity, and overall variation), which we used for comparison across native and QTY analogs. We also used its predictions of transmembrane helix topology to compare the topology of native ATP synthase Fo subunits and their corresponding QTY analogs.

### Structural Models

When available, native structures were retrieved from experimentally determined CryoEM models available in the RCSB <u>P</u>rotein <u>D</u>ata <u>B</u>ank (RCSB PDB), specifically state 1 of the human ATP synthase CryoEM structure (PDB 8H9S) (https://www.rcsb.org/structure/8H9S/), which was chosen for its singular and well-defined conformational structure. For proteins without an experimentally determined PDB structure, native models were generated via AlphaFold3 servers (https://alphafoldserver.com/) from corresponding UniProt sequences. QTY analogs were generated in the same way, using QTY-designed sequences as inputs for AlphaFold3 predictions. AlphaFold3 output visualizations were captured as screenshots for analysis preparation.

### Rationale for Separating Comparisons into Native CryoEM and AlphaFold3-Predicted Native Structures

We separated structural comparisons into native CryoEM and native AlphaFold3 categories because experimental CryoEM structures were not available for all ATP synthase Fo subunits. ATP6, ATP8, ATPK, and ATP68 each had PDB entries with CryoEM structures, allowing us to compare their native CryoEM structures with their QTY analogs.

Meanwhile, ATPMK, AT5G1, AT5G2, and AT5G3 were analyzed using AlphaFold3-predicted native structures. Although AT5G1 had a PDB entry, its CryoEM structure was incomplete and lacked amino acids. To avoid bias from missing residues, we used AlphaFold3 to generate the full native structure to compare with its QTY analog. We did the same with AT5G2, AT5G3, and ATPMK, which all lacked PDB entries. This separation was made to ensure that each QTY analog was compared against the most complete and appropriate native structure.

### Structural Superposition and RMSD

Native and QTY protein structures were superposed in PyMOL (https://pymol.org), and RMSDs were computed from the resulting alignments. We used unrelaxed RMSD values for all reporting. PyMOL (https://pymol.org) was used to superpose the native protein structure and the QTY variant, and calculate the RMSDs. For simplicity and clarity, unstructured loops and extraneous protein monomers were removed from the figures before superposition.

### Structure Visualization

PyMOL (https://pymol.org) was used to render the superpositions. ATP68 was divided into two segments and connected in PyMOL because the CryoEM structure showed a clear bend, whereas the AlphaFold3 version did not. Segmenting the protein reduced the misalignment caused by this conformation difference and allowed for clearer visual comparison. UCSF ChimeraX (https://www.rbvi.ucsf.edu/chimerax) was used to render protein models with hydrophobicity patches. Transmembrane

### Data availability of AlphaFold-predicted water-soluble QTY analogs

The QTY code designed water-soluble analogs of the human integral membrane protein enzymes are available at https://github.com/timwang08/Human-ATP-Synthase-Fo-QTY-Analogs. If additional information is needed, please contact Z.W., Protein characteristics used in the analysis are available on UniProt (https://www.uniprot.org/). The native CryoEM-determined four ATP synthase Fo subunits are available in the RCSB PDB repository (https://www.rcsb.org/).

